# Mutagenesis-Visualization: Analysis of Site-Saturation Mutagenesis Datasets in Python

**DOI:** 10.1101/2021.10.08.463725

**Authors:** Frank Hidalgo, Sage Templeton, Che Olavarria Gallegos, Joanne Wang

## Abstract

**Summary:** Site-saturation mutagenesis experiments have been transformative in our study of protein function. Despite the rich data generated from such experiments, current tools for processing, analyzing, and visualizing the data offer only a limited set of static visualization tools that are difficult to customize. Furthermore, usage of the tools requires extensive experience and programming knowledge, slowing the research process for those in the biological field who are unfamiliar with programming. Here, we introduce *mutagenesis-visualization*, a Python package for creating publication-quality figures for site-saturation mutagenesis datasets without the need for prior Python or statistics experience, where each of the graphs is generated with a one-line command. The plots can be rendered as native *Matplotlib* objects (easy to stylize) or *Plotly* objects (interactive graphs). Additionally, the software offers the possibility to visualize the datasets on *Pymol*.

**Availability and implementation:** The software can be installed from *PyPI* or *GitHub* using the *pip* package manager and is compatible with Python ≥ 3.8. The [documentation](https://mutagenesis-visualization.readthedocs.io/) can be found at *readthedocs* and the [source code](https://github.com/fhidalgor/mutagenesis_visualization) on *GitHub*.

## Introduction

Site-saturation mutagenesis (also known as deep mutational scanning) allows for the simultaneous, unbiased, and comprehensive study of all possible amino acid substitutions for a protein (Tripathi and Varadarajan 2014; Fowler and Fields 2014; Nov 2012; Shin and Cho 2015; Wrenbeck et al. 2017; Araya and Fowler 2011). First, a library of single-codon variants is plugged into an assay, where mutations in the protein affect functional selection. Next, the library is retrieved from the before and after selection samples. Then, using next-generation sequencing, the counts of each variant are used to calculate an enrichment score. Numerous applications have been pursued using site-saturation mutagenesis, including drug resistance prediction (Pines et al. 2020), protein engineering (Shin and Cho 2015), allostery determination (Subramanian et al. 2021), and functional analysis of genomes (Li et al. 2016).

A substantial number of software tools for different parts of the bioinformatics pipeline have been described in the literature. For example, for DNA library design, *Mutation Maker* and *OneClick* let users design oligos according to different specifications (Tang et al. 2012; Hiraga et al. 2021). For the pre-processing of DNA reads, *FLASH* and *PEAR* tools merge paired-end reads, *Cutadapt* and *Trimmomatic* remove adapter sequences (Zhang et al. 2014; Bolger et al. 2014), and QUASR provides a framework for quantification and analysis of short reads (Gaidatzis et al. 2015). To calculate enrichment scores and statistical analysis, *DiMSsum*, *enrich2*, *dms*_*tools2,* and *DESeq2* use different statistical models to quantify the errors and treat replicates and time series (Love et al. 2014; Rubin et al. 2017; Bloom 2015; Faure et al. 2020). This last subset of tools may also integrate a pre-processing module. For identifying molecular constraints that affect enrichment scores, there is *dms2dfe* (Dandage and Chakraborty 2017). Lastly, the Bloom lab recently released *dms-view*, a web-based JavaScript tool that lets users map site-saturation mutagenesis datasets to a 3-D protein structure (Hilton et al. 2020).

While the current tools allow users to calculate enrichment scores and error estimates from raw sequencing data, they only have a few built-in static visualization options such as heatmaps, logos, and scatter plots. Customizing these plots, or creating different plots, requires a high knowledge of programming and extensive data manipulation, making the creation of publication-quality figures challenging and time-consuming. In addition, none of these packages allows plots to be integrated with *Pymol* (Schrödinger 2015), the go-to molecular visualization tool for scientists. Thus, overlaying enrichment scores onto a PDB structure must be done by hand, slowing the data analysis process. Here, we describe *mutagenesis-visualization*, a Python package that addresses the aforementioned needs.

## Implementation

*Mutagenesis-visualization* is a user-centered Python package for processing, analyzing, and creating high-quality figures for publication from pairwise site-saturation mutagenesis datasets. Unlike other packages, *mutagenesis-visualization* handles all the data manipulation internally, streamlining the workflow. The software workflow consists of two main steps: (i) data processing (count of DNA reads and calculation of enrichment scores) and (ii) data analysis and visualization. The first step counts the DNA reads for each variant, and then, from pre-selection and post-selection samples, calculates the enrichment score for each variant. *Mutagenesis-visualization* has several options for processing and normalizing datasets. Due to the modularity of our software, this first step can be conducted separately on other available software packages used to process site-saturation mutagenesis data, and the data can be integrated back into the *mutagenesis-visualization* pipeline for further visualization analysis. Once the enrichment scores are calculated, different types of visualizations can be conducted.

These datasets are commonly represented with heatmaps. In *mutagenesis-visualization*, heatmaps are highly tunable (Figure 1A). The user can change all stylistic aspects such as size, labels, color scheme, and protein cartoon. These heatmaps can also be customized by choosing protein residues of interest or by selecting a subset of amino acids. Furthermore, heatmaps with average substitutions can be generated to easily visualize global phenotypic trends of the variants (Figure 1B). Since heatmaps are just one method of visualizing data, *mutagenesis-visualization* provides tools for plotting data sets through histograms, scatter plots, line plots, and bar charts (Figure 1D, 1E, 1H). Each of the graphs shown in Figure 1 is generated with a one-line command.

**Figure 1.**
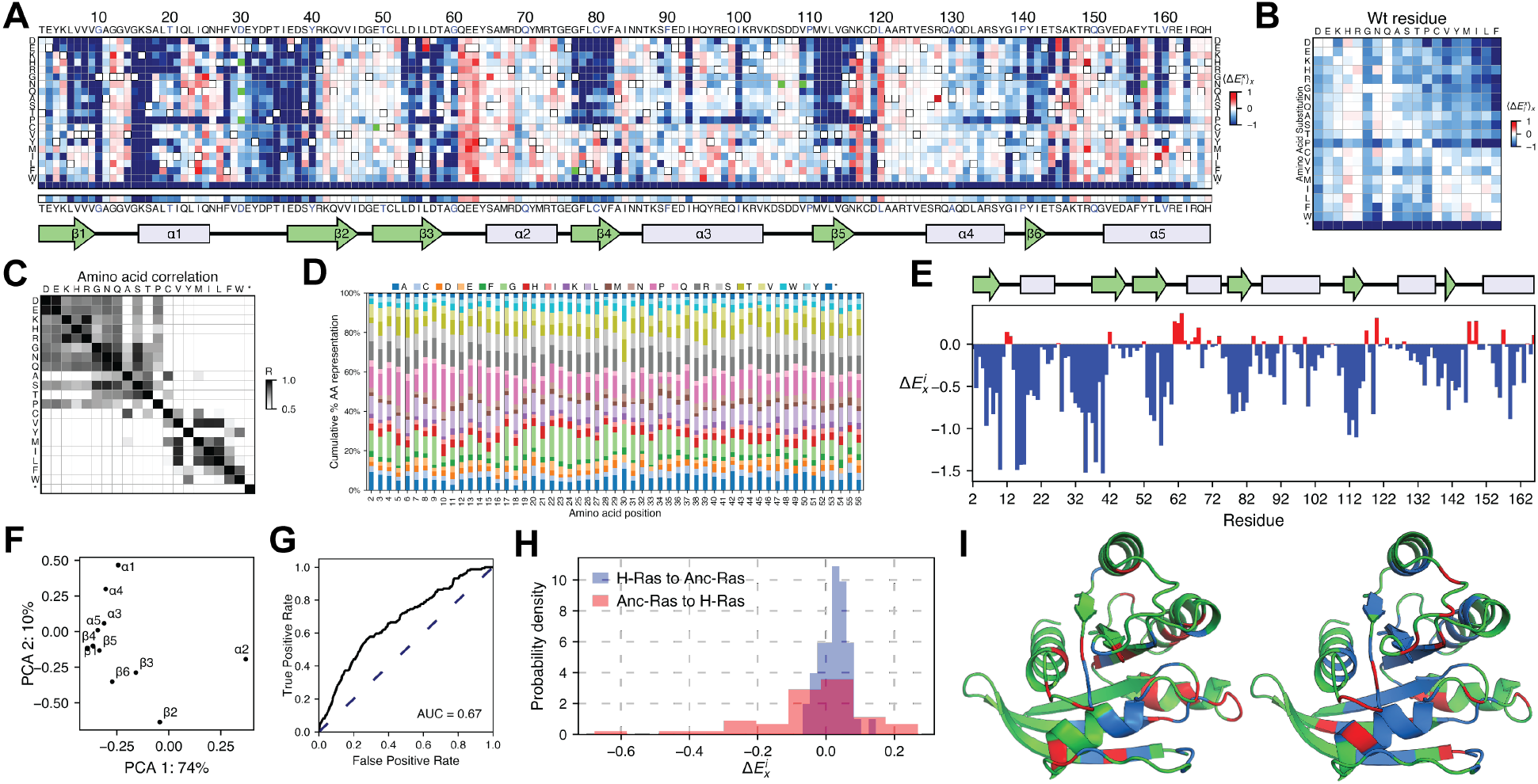
Example of *mutagenesis-visualization* types of plots. The user passes into the software a *pandas* dataframe (McKinney 2010) or *NumPy* array (Harris et al. 2020) containing the enrichment scores, the protein sequence, and, optionally, the secondary structure. The user can generate each of the following plots using a one-line command. The data used to generate the plots was obtained from replicating previous work on H-Ras (Bandaru et al. 2017). For figures A, B, E, and I, shades of red and blue indicate gain and loss of function, respectively. (**A**) A heatmap representing enrichment scores for substituting a particular residue in H-Ras with one of the 20 amino acids. The cartoon indicates the secondary structure motifs. (**B**) A heatmap with average amino acid substitution effects in H-Ras. (**C**) A correlation plot where the Pearson R-value is calculated for each amino acid substitution. (**D**) A bar plot representing the cumulative percentage of each amino acid substitution in the DNA library grouped by residue. The library was used to conduct the H-Ras site-saturation mutagenesis experiment. Each amino acid is coded with a different color. (**E**) A bar plot representing the average amino acid substitution effect per residue in H-Ras. (**F**) A scatter plot of the first two dimensions of a principal component analysis (PCA) conducted on the H-Ras dataset, which had been grouped by the secondary structure elements. (**G**) A receiver operating characteristic (ROC) plot used to make a quantitative determination of the fit between the measured enrichment scores and binary classification of the variants. The binary classification could represent any property such as sequence conservation, whether found in a cancer screen database or any other categorization that the user wants. (**H**) A histogram plot of the enrichment scores of all the sequence changes required for the H-Ras protein sequence to revert to Ancestral Ras and vice versa. There are forty-eight residues that differ between the two protein sequences (Bandaru et al. 2017). The data from the Ancestral Ras experiment are unpublished. (**I**) Two *Pymol*-generated figures where the alanine (left) and aspartate (right) enrichment scores were mapped onto the Ras structure (PDB ID: 5P21) on *Pymol*. The user needs a valid *Pymol* license and the same Python path for *Pymol* and for the virtual environment used to run this software.

Moreover, *mutagenesis-visualization* includes tools for hierarchical clustering and correlations (Figure 1C), PCA (Figure 1G), and ROC curves (Figure 1F). In addition, the *matplotlib* axes objects (Hunter 2007) can be retrieved, allowing for a high level of customization. Moreover, most visualizations, including heatmaps, can also be generated using *Plotly*, allowing interactive plots with hover features and dashboard creation. We have created a sample [dashboard](http://mvhras.herokuapp.com/) on *Heroku* to illustrate the capabilities of the software. Lastly, *mutagenesis-visualization* allows data to be easily exported and visualized onto a PDB structure in *Pymol* (Figure 1I).

*Mutagenesis-visualization* is designed to be run locally and does not require a high-performance computing environment. The software can be installed from *PyPI* or *GitHub* using the *pip* package manager and is compatible with Python ≥ 3.8. The source code is available in the *GitHub* [repository](https://github.com/fhidalgor/mutagenesis_visualization). The [documentation](http://mutagenesis-visualization.readthedocs.io/) contains a step-by-step tutorial on the different types of plots, and a live version of the [tutorial](https://mybinder.org/v2/gh/fhidalgor/mutagenesis_visualization/HEAD?filepath=mutagenesis_visualization/tutorial/) is hosted on *mybinder*, and can be run remotely without any installation. In addition, the tutorial includes the analysis of sample site-saturation mutagenesis datasets from various studies (Dou et al. 2018; Fernandes et al. 2016; Livesey and Marsh 2020; Melnikov et al. 2014; Newberry et al. 2020; Stiffler et al. 2015; Bandaru et al. 2017).

## Conclusion

*Mutagenesis-visualization* fills a need in the Python and scientific community for user-friendly software that streamlines the pairwise site-saturation mutagenesis bioinformatics pipeline, where each of the graphs can be generated with a one-line command. While the workflow allows for end-to-end analysis of the pre-processed datasets, its modularity makes it compatible with other available software used to process site-saturation mutagenesis data. [Source code](https://github.com/fhidalgor/mutagenesis_visualization), [documentation](https://mutagenesis-visualization.readthedocs.io/), as well as a comprehensive, step-by-step [tutorial](https://mybinder.org/v2/gh/fhidalgor/mutagenesis_visualization/HEAD?filepath=mutagenesis_visualization/tutorial/) can be found online.

## Acknowledgments

The authors thank Prof. John Kuriyan, Subu Subramanian, and all the members of the Kuriyan lab for their insightful discussions.

## Bibliography

Araya, C.L. and Fowler, D.M. 2011. Deep mutational scanning: assessing protein function on a massive scale. Trends in Biotechnology 29(9), pp. 435–442.

Bandaru, P., Shah, N.H., Bhattacharyya, M., et al. 2017. Deconstruction of the Ras switching cycle through saturation mutagenesis. eLife 6.

Bloom, J.D. 2015. Software for the analysis and visualization of deep mutational scanning data. BMC Bioinformatics 16, p. 168.

Bolger, A.M., Lohse, M. and Usadel, B. 2014. Trimmomatic: a flexible trimmer for Illumina sequence data. Bioinformatics 30(15), pp. 2114–2120.

Dandage, R. and Chakraborty, K. 2017. dms2dfe: Comprehensive Workflow for Analysis of Deep Mutational Scanning Data. The Journal of Open Source Software 2(20), p. 362.

Dou, J., Vorobieva, A., Sheffler, W., et al. 2018. De Novo Design Of A Fluorescence-Activating Β-Barrel. Zenodo.

Faure, A.J., Schmiedel, J.M., Baeza-Centurion, P. and Lehner, B. 2020. DiMSum: an error model and pipeline for analyzing deep mutational scanning data and diagnosing common experimental pathologies. Genome Biology 21(1), p. 207.

Fernandes, J.D., Faust, T.B., Strauli, N.B., et al. 2016. Functional segregation of overlapping genes in HIV. Cell 167(7), p. 1762–1773.e12.

Fowler, D.M. and Fields, S. 2014. Deep mutational scanning: a new style of protein science. Nature Methods 11(8), pp. 801–807.

Gaidatzis, D., Lerch, A., Hahne, F. and Stadler, M.B. 2015. QuasR: quantification and annotation of short reads in R. Bioinformatics 31(7), pp. 1130–1132.

Harris, C.R., Millman, K.J., van der Walt, S.J., et al. 2020. Array programming with NumPy. Nature 585(7825), pp. 357–362.

Hilton, S.K., Huddleston, J., Black, A., et al. 2020. dms-view: Interactive visualization tool for deep mutational scanning data. Journal of open source software 5(52).

Hiraga, K., Mejzlik, P., Marcisin, M., et al. 2021. Mutation maker, an open source oligo design platform for protein engineering. ACS synthetic biology [electronic resource] 10(2), pp. 357–370.

Hunter, J.D. 2007. Matplotlib: A 2D Graphics Environment. Computing in science & engineering 9(3), pp. 90–95.

Li, C., Qian, W., Maclean, C.J. and Zhang, J. 2016. The fitness landscape of a tRNA gene. Science 352(6287), pp. 837–840.

Livesey, B.J. and Marsh, J.A. 2020. Using deep mutational scanning to benchmark variant effect predictors and identify disease mutations. Molecular Systems Biology 16(7), p. e9380.

Love, M.I., Huber, W. and Anders, S. 2014. Moderated estimation of fold change and dispersion for RNA-seq data with DESeq2. Genome Biology 15(12), p. 550.

McKinney, W. 2010. Data structures for statistical computing in python. In: Proceedings of the 9th Python in Science Conference. Proceedings of the python in science conference. SciPy, pp. 56–61.

Melnikov, A., Rogov, P., Wang, L., Gnirke, A. and Mikkelsen, T.S. 2014. Comprehensive mutational scanning of a kinase in vivo reveals substrate-dependent fitness landscapes. Nucleic Acids Research 42(14), p. e112.

Newberry, R.W., Leong, J.T., Chow, E.D., Kampmann, M. and DeGrado, W.F. 2020. Deep mutational scanning reveals the structural basis for α-synuclein activity. Nature Chemical Biology 16(6), pp. 653–659.

Nov, Y. 2012. When second best is good enough: another probabilistic look at saturation mutagenesis. Applied and Environmental Microbiology 78(1), pp. 258–262.

Pines, G., Fankhauser, R.G. and Eckert, C.A. 2020. Predicting drug resistance using deep mutational scanning. Molecules (Basel, Switzerland) 25(9).

Rubin, A.F., Gelman, H., Lucas, N., et al. 2017. A statistical framework for analyzing deep mutational scanning data. Genome Biology 18(1), p. 150.

Schrödinger, L.L.C. 2015. The {PyMOL} Molecular Graphics System, Version~1.8.

Shin, H. and Cho, B.-K. 2015. Rational protein engineering guided by deep mutational scanning. International Journal of Molecular Sciences 16(9), pp. 23094–23110.

Stiffler, M.A., Hekstra, D.R. and Ranganathan, R. 2015. Evolvability as a function of purifying selection in TEM-1 β-lactamase. Cell 160(5), pp. 882–892.

Subramanian, S., Gorday, K., Marcus, K., et al. 2021. Allosteric communication in DNA polymerase clamp loaders relies on a critical hydrogen-bonded junction. eLife 10.

Tang, L., Gao, H., Zhu, X., Wang, X., Zhou, M. and Jiang, R. 2012. Construction of “small-intelligent” focused mutagenesis libraries using well-designed combinatorial degenerate primers. Biotechniques 52(3), pp. 149–158.

Tripathi, A. and Varadarajan, R. 2014. Residue specific contributions to stability and activity inferred from saturation mutagenesis and deep sequencing. Current Opinion in Structural Biology 24, pp. 63–71.

Wrenbeck, E.E., Faber, M.S. and Whitehead, T.A. 2017. Deep sequencing methods for protein engineering and design. Current Opinion in Structural Biology 45, pp. 36–44.

Zhang, J., Kobert, K., Flouri, T. and Stamatakis, A. 2014. PEAR: a fast and accurate Illumina Paired-End reAd mergeR. Bioinformatics 30(5), pp. 614–620.

